# MR-DoC2: Bidirectional Causal Modeling with Instrumental Variables and Data from Relatives

**DOI:** 10.1101/2022.03.14.484271

**Authors:** Luis FS Castro-de-Araujo, Madhurbain Singh, Yi (Daniel) Zhou, Philip Vinh, Brad Verhulst, Conor V Dolan, Michael C Neale

## Abstract

Establishing causality is an essential step towards developing interventions for psychiatric disorders, substance use and many other conditions. While randomized controlled trials (RCTs) are considered the gold standard for causal inference, they are unethical in many scenarios. Mendelian randomization (MR) can be used in such cases, but importantly both RCTs and MR assume unidirectional causality. In this paper, we developed a new model, MRDoC2, that can be used to identify bidirectional causation in the presence of confounding due to both familial and non-familial sources. Our model extends the MRDoC model (Minică et al 2018), by simultaneously including risk scores for each trait. Furthermore, the power to detect causal effects in MRDoC2 does not require the phenotypes to have different additive genetic or shared environmental sources of variance, as is the case in the direction of causation twin model (Heath et al., 1993).

## Introduction

To understand the mechanisms of diseases and disorders, it is essential to differentiate between correlation and causation. Many research designs and current statistical models are unable to distinguish correlation from causation. While randomized controlled trials (RCTs) are the gold standard for causal inference (Evans and Davey Smith, 2015), they are unethical or impractical in many common situations. For example, exposure to a traumatic experience, a possible risk factor for substance abuse, is not amenable to randomization. Mendelian randomization (MR), an instrumental variable method, is an alternative approach to evaluate causality when RCTs are not possible. It has improved our understanding of the etiologies of several conditions (Ohlsson and Kendler, 2020). However, MR is based on strong assumptions, which can be difficult to validate. In this article, we consider research designs that mitigate such assumptions.

MR is an important method for examining causality in observational studies (Choi et al., 2020; Katikireddi et al., 2018). It uses genetic variants as instrumental variables to detect and estimate the causal effect of an exposure on an outcome. This method has three main assumptions: i) exchangeability, the variant is not associated with a confounder in the relation between the exposure and outcome; ii) no horizontal pleiotropy (exclusion restriction), the variant should affect the outcome exclusively via the exposure; and iii) relevance, the instruments are sufficiently predictive of the exposure. Most psychiatric and substance use disorders appear to be multifactorial and highly polygenic, i.e., they are subject to the effects of many environmental events and genetic variants (e.g., SNPs) with small effects on the phenotype of interest. As instruments, individual genetic variants are likely to be poorly predictive of the exposure, which renders MR susceptible to weak instrument bias. The bias can be in the direction of the confounded observed association, as previously reported (Burgess et al., 2011; Burgess and Thompson, 2013; Hemani et al., 2018). The size and direction of the bias, however, depends on the research study design.

The instrument should not share common causes with the outcome and should affect the outcome exclusively via the exposure, which is known as the *no horizontal pleiotropy* assumption. The *no horizontal pleiotropy* assumption seems unlikely to be met in complex traits such as psychiatric disorders or substance use, because the relevant phenotypes and variants are characterized by pervasive comorbidity and pleiotropy. With larger genome-wide association studies (GWAS), more variants are found to have pleiotropic effects, with hundreds being reported (Jordan et al., 2019). In two-sample MR, the data originate from two separate GWAS, so no individual has data on both exposure and outcome. By contrast, in one-sample MR, genotyped individuals are assessed on both traits. In one-sample MR, weak instrument bias tends to be in the direction of the confounded observed association, whereas in two-sample MR, the bias is towards the null (i.e., regression dilution; Hemani et al., 2018).

In MR, the causation is assumed to be unidirectional from exposure to outcome (Hemani et al., 2018; Katikireddi et al., 2018). Some authors have applied MR to investigate directionality of the causal paths (Timpson et al., 2011; Welsh et al., 2010). However, here directionality is investigated by performing MR twice, one test for each direction between the variables of interest. This can be problematic, as the relationship in one direction can interfere with the relationship in the opposite direction. Biology features many such feedback loops of this kind, from homeostatic mechanisms for body temperature or nicotine level (Verhulst et al 2021) to the mutual potentiation of aggressive behavior and punitive discipline (Smith, 2006).

Like RCTs, MR can be applied to infer causality in the presence of confounding, but the approach comes with assumptions that limit its application in some cases. Other designs can be helpful in some cases where MR is not suitable given its assumptions. Here we explored a structural equation model (SEM) implementation of MR, which allows for full background confounding and reciprocal causation. This model combines two twin models, namely the Direction of Causation model and the MR-DoC model (Minica et al, 2018). The Direction of Causation (DoC) model can infer causal relationships by using information from the cross-twin cross-trait correlations, even in cross-sectional studies (Neale & Cardon 1992; Heath et al 1993; Neale et al., 1994). This model requires two phenotypes, which differ in their MZ correlations, or their DZ correlations, or both. However, to accommodate bidirectional causality, one must assume that confounding factors are limited to either additive genetic, common environment or specific environmental components of variance. In essence, any three of the five potential sources of covariance between the traits may be estimated, but no more (Figure 1; Duffy and Martin, 1994; Maes et al., 2021).

**Figure 1.**
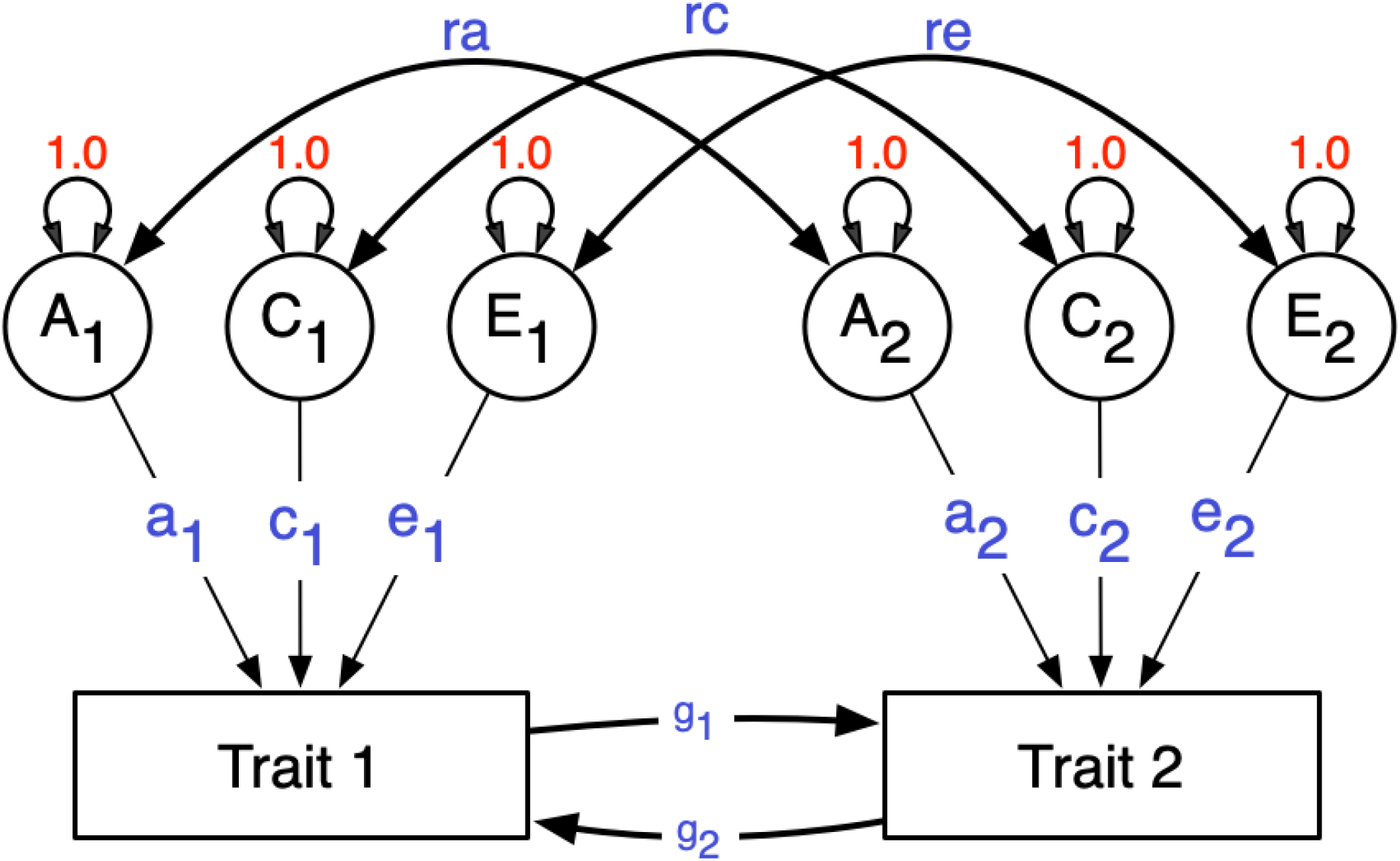
Classic DoC model. Path diagram representing a Direction of Causation model for one twin. This is depicting the relationship between two phenotypes, and the causal paths are estimated affording information from the cross-twin cross-trait correlations. Cross-twin covariance between additive genetic effects is 0.5 (not shown) for DZ twins, as DZs are expected to share 50% of the genetic effects. Standard structural equation modeling symbology is used. Circles represent latent variables, whose variances are fixed to unity. Double-headed arrows are covariances or variances, single-headed are the causal regression paths. Squares represent observed variables. A, C and E are the additive, shared and unique environmental effects. This model is not identified with data from MZ and DZ twins. To make it identified one needs to set to zero any two of the five paths *ra, rc, re, g_2_* and *g_1_* (Maes et al., 2021).

Minică et al. (2018) combined the classical twin model and the MR model (Heath et al 1993; Gillespie et al., 2003; Neale et al., 1994), in a model for unidirectional causality called MR-DoC. This model accommodates horizontal pleiotropy, i.e., an effect of the genetic instrument on the outcome that is not mediated by the exposure variable. These alternative pathways are often denoted “direct effects,” but in practice the alternative pathways likely involve many different mediators. Following Burgess et al. (2011), Burgess and Thompson (2013), and Kohler et al. (2011), Minică et al. (2018) used a polygenic score (PS) as the instrument, in an attempt to avoid the weak instrument problem associated with single SNPs. Minică et al. (2018) showed that the model has higher power to detect causality than traditional MR (Minică et al., 2018). Two strong assumptions of the MR-DoC model are unidirectional causation and the absence of within-person (i.e., unshared) environmental confounding. The latter implies that the correlation between the non-familial environmental influences on the exposure and the outcome is zero. In contrast, confounding originating in genetic and shared environmental correlations can be accommodated in the MR-DoC model.

MR-DoC2 extends the MR-DoC model by integrating two polygenic scores (PSs), one for each phenotype. Using two PSs in a twin sample allows us to model bidirectional causation in the presence of full confounding, originating in shared and unshared environmental and additive genetic effects. The ability to estimate bidirectional causality in the presence of full (shared and unshared environmental and additive genetic) confounding requires the assumption of no direct horizontal pleiotropy: we assume no direct effect of PS1 on phenotype 2 (Ph2), nor of PS2 on phenotype 1 (Ph1). However, we note that the model does not imply that PS1 (or PS2) is uncorrelated with Ph2 (Ph1), we return to this issue below. We first present the model, address the issue of parameter identification, and continue to address statistical power.

## Methods

All analyses were performed using R version 4.1.1 (R Core Team, 2021). The models were specified using RAM matrix algebra in OpenMx v2.19.6 (Neale et al., 2016). We established local model identification numerically using the OpenMx function mxCheckIdentification (Hunter et al., 2021). The MR-DoC 2 model is shown in Figure 2 for an individual twin (rather than a twin pair) to ease presentation. In Figure 2, the two phenotypes (denoted Ph1 and Ph2) are modeled as follows:

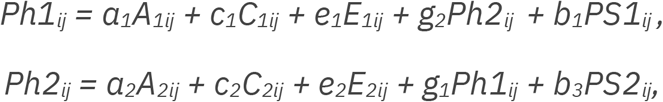

where *i* denotes twin pair and *j* denotes individual twin. The parameters *a_1_, c_1_* and *e_1_* represent the effects of the additive genetic (*A*), shared environmental (*C*), and unshared environmental variables (*E*) on the phenotype *Ph1 (a_2_, c_2_*, and *e_2_*, defined analogously as effects of *A_2_, C_2_*, and *E_2_ on Ph_2_*). The parameters *b_1_* and *b_3_* express the instrumental variable effects on *Ph1* and *Ph2*, respectively. The parameters of main interest are *g_1_* and *g_2_*, i.e., the bidirectional causal effects. Note that the model includes correlations between the additive genetic and environmental variables (i.e., cor(*A_1_, A_2_*)=*ra*, etc.), which are denoted *ra, rc*, and *re*. An important feature of the present model is that it includes the parameter *re*, as it has been previously shown that the absence of *re* in a DoC model would introduce bias in the estimates (Rasmussen et al., 2019). Finally, note that the instruments may be correlated (correlation *rf* in Figure 2), due to possible linkage disequilibrium.

**Figure 2.**
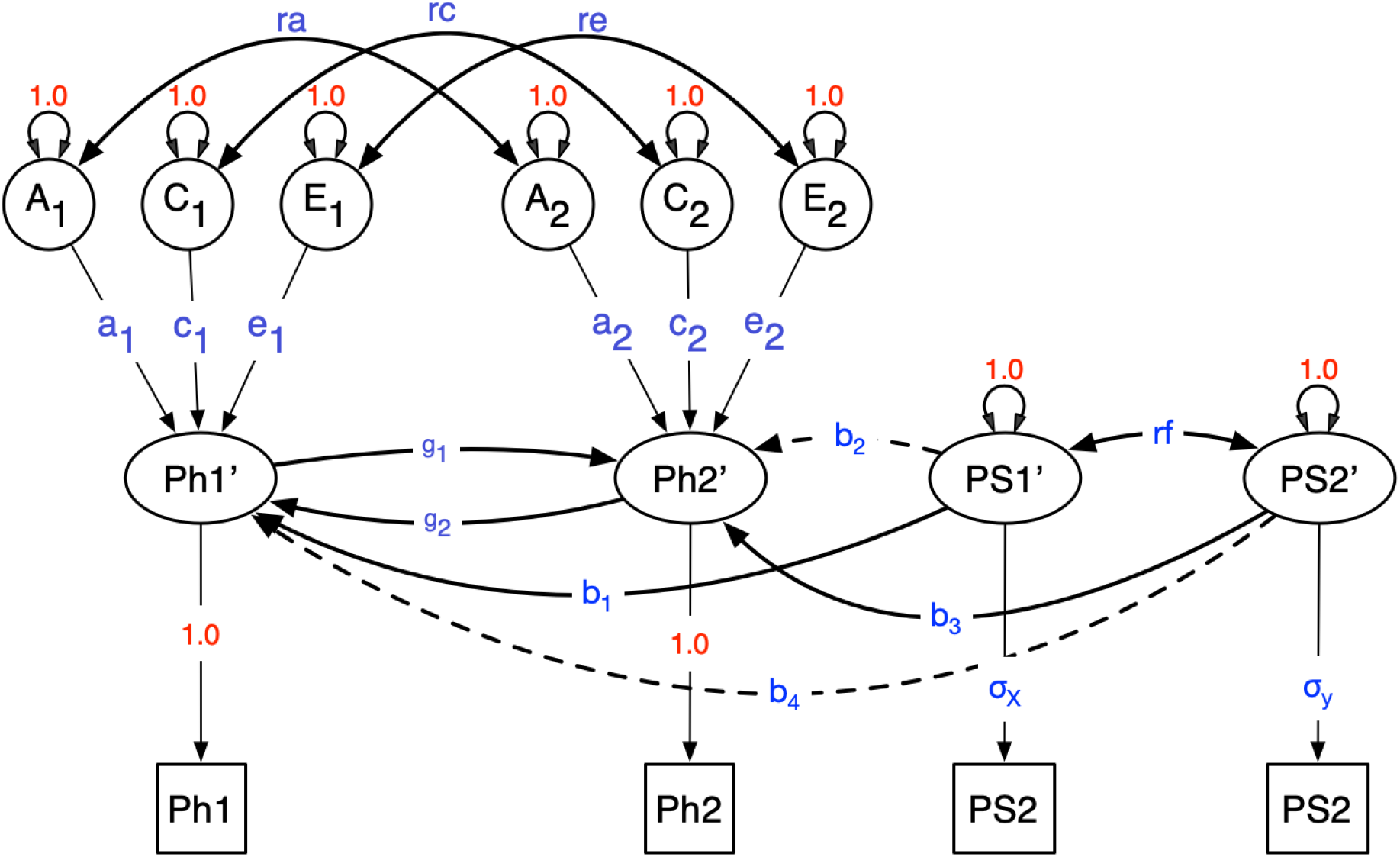
MR-DoC2. Path diagram of the MR-DoC2 model for an individual. The model is locally identified when parameters *b_2_* and *b_4_* are fixed (zero). The model includes the effects of additive genetic (*A*), common environment (*C*) and specific environment (*E*) factors for both *Ph1* and *Ph2*, and their effects may correlate (parameters *ra, rc* and *re*).

The vector of parameters is denoted θ = [*ra, rc, re, rf, a-_1_, c_1_, -*e*_1_, α_2_, c_2_, -e_2_, -g_2_, -g_2_, b_1_, b_3_, σ_x_, σ_y_*], where σ_*x*_ and σ_*y*_ are the standard deviations of the PS1 and PS2, respectively. The model’s path coefficients and variances were estimated by maximum likelihood estimation; see details of the simulation next section (Tables 2, 3, and S1). Note that the model, as shown in Figure 2, includes the parameters *b_2_* and *b_4_*, but these parameters were fixed to zero for identification. However, possible combinations of parameters that render the model identified are presented in Table 5. In particular, one cannot freely estimate *b_2_* and *b_4_*, but either can be estimated if *re* is fixed to zero (Table 5).

**Table 1.** Parameter levels on the three factorial designs, with respective total number of cells for each design simulation. The model specification can be seen in Figure 2. Also, *e_1_* was specified as 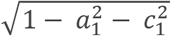 and *e_2_* as 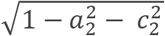. Table 4 exemplifies rows extracted from Design 3.

| $\theta$ | Design 1 (ACE) | Design 2 (AE) | Design 3 (AE) |
| --- | --- | --- | --- |
| $b_1$ | $\sqrt{0.025}, \sqrt{0.05}$ | $\sqrt{0.025}, \sqrt{0.05}, \sqrt{0.075}$ | $-\sqrt{0.075}, -\sqrt{0.03}, \sqrt{0.03}, \sqrt{0.075}$ |
| $b_3$ | $\sqrt{0.025}, \sqrt{0.05}$ | $\sqrt{0.025}, \sqrt{0.05}, \sqrt{0.075}$ | $-\sqrt{0.075}, -\sqrt{0.03}, \sqrt{0.03}, \sqrt{0.075}$ |
| $g_1$ | $\sqrt{0.020}, \sqrt{0.05}$ | $\sqrt{0.020}, \sqrt{0.04}, \sqrt{0.06}$ | $-\sqrt{0.050}, -\sqrt{0.020}, \sqrt{0.050}, \sqrt{0.020}$ |
| $g_2$ | $\sqrt{0.020}, \sqrt{0.05}$ | $\sqrt{0.020}, \sqrt{0.04}, \sqrt{0.06}$ | $-\sqrt{0.050}, -\sqrt{0.020}, \sqrt{0.050}, \sqrt{0.020}$ |
| $ra$ | $0.25, 0.50$ | $0.0, 0.25, 0.50$ | 0.3 |
| $rc$ | $0.25, 0.50$ | 0 | 0 |
| $re$ | $0.25, 0.50$ | $0.0, 0.25, 0.50$ | 0.3 |
| $rf$ | $0.25, 0.50$ | $0.0, 0.25, 0.50$ | 0.3 |
| $a_1$ | $\sqrt{0.10}, \sqrt{0.25}$ | $\sqrt{0.10}, \sqrt{0.25}$ | $\sqrt{0.5}$ |
| $a_2$ | $\sqrt{0.10}, \sqrt{0.25}$ | $\sqrt{0.10}, \sqrt{0.25}$ | $\sqrt{0.3}$ |
| $c_1$ | $\sqrt{0.10}, \sqrt{0.25}$ | 0 | 0 |
| $c_2$ | $\sqrt{0.10}, \sqrt{0.25}$ | 0 | 0 |
| **Total cells** | $2^{12}=4096$ | $3^7 \cdot 2^2= 8748$ | $4^4=256$ |

**Table 2.** Variance explained in statistical power (non-centrality parameter; NCP) by model parameters

| | NCP ($\beta^2$) | | |
| --- | --- | --- | --- |
| | $g_1 = 0$ | $g_2 = 0$ | $g_1 = g_2 = 0$ |
| **b<sub>1</sub>** | 0.365 | 0.000 | 0.181 |
| **g<sub>1</sub>** | 0.517 | 0.000 | 0.289 |
| **b<sub>3</sub>** | 0.000 | 0.365 | 0.181 |
| **g<sub>2</sub>** | 0.000 | 0.517 | 0.289 |
| **ra** | 0.000 | 0.000 | 0.000 |
| **rc** | 0.000 | 0.000 | 0.000 |
| **re** | 0.002 | 0.002 | 0.000 |
| **rf** | 0.041 | 0.041 | 0.000 |
| **a<sub>1</sub>** | 0.000 | 0.002 | 0.001 |
| **a<sub>2</sub>** | 0.002 | 0.000 | 0.001 |
| **c<sub>1</sub>** | 0.000 | 0.002 | 0.001 |
| **c<sub>2</sub>** | 0.002 | 0.000 | 0.001 |
| **Total R<sup>2</sup>** | 0.929 | 0.929 | 0.945 |

Note: Linear regression of NCP on the parameter values used for 4096 power analyses in the ACE model.

**Table 3.** Variance explained in statistical power (non-centrality parameter; NCP) by model parameters

| | NCP ($\beta^2$) | | |
| --- | --- | --- | --- |
| | $g_1 = 0$ | $g_2 = 0$ | $g_1 = g_2 = 0$ |
| **b<sub>1</sub>** | 0.460 | 0.000 | 0.226 |
| **g<sub>1</sub>** | 0.402 | 0.000 | 0.222 |
| **b<sub>3</sub>** | 0.000 | 0.460 | 0.226 |
| **g<sub>2</sub>** | 0.000 | 0.402 | 0.222 |
| **ra** | 0.001 | 0.001 | 0.001 |
| **rc** | 0.000 | 0.000 | 0.000 |
| **re** | 0.009 | 0.009 | 0.001 |
| **rf** | 0.036 | 0.036 | 0.001 |
| **a<sub>1</sub>** | 0.000 | 0.003 | 0.002 |
| **a<sub>2</sub>** | 0.003 | 0.000 | 0.002 |
| **c<sub>1</sub>** | 0.000 | 0.000 | 0.000 |
| **c<sub>2</sub>** | 0.000 | 0.000 | 0.000 |
| **Total R<sup>2</sup>** | 0.911 | 0.911 | 0.902 |

Note: Linear regression of NCP on the parameter values used for 8748 power analyses in the AE model.

### Simulation procedure

Statistical power was explored by exact data simulation given various combinations of parameter values (van der Sluis et al., 2008). We used the R function mvrnorm() in the MASS library to simulate exact data (Venables et al., 2002). The power calculation was based on the likelihood ratio (LR) test. Given exact data simulation, parameter estimates of the full (identified) model equal the true values exactly. By subsequently fitting a model with one or more parameters of choice fixed to zero (nested under the true model), we obtain the exact non-centrality parameter (NCP) of the LR test given the sample sizes. The NCP is the difference in the expected value of the LR test statistic under the null and the alternative hypotheses (Verhulst, 2017). Given the NCP, the degrees of freedom of the LT test (i.e., the difference in the number of parameters in the null and the alternative hypothesis), and the chosen Type I error rate alpha (e.g., 0.05), we can calculate the power to reject the constraints associated with the alternative hypothesis. The exact data simulation approach is equivalent to the analysis of exact population covariance matrices and mean vectors. While power can be established empirically using simulated data, this is computationally less efficient and offers no advantage above exact data simulation.

The simulation procedure involved: 1) choosing a set of parameter values for the model shown in Figure 2; 2) exact data simulation, with arbitrary N=1,000 MZ pairs and N=1,000 DZ twin pairs; 3) fitting the true model using ML estimation in OpenMx; 4) fitting the false model by fixing one or more parameters to zero and refitting the model; and 5) calculating the NCP and the power to reject the false model restrictions. Type I error rate α was set to 0.05. In the power calculations, we focused on the causal parameters *g_1_*, or *g_2_*, or *g_1_* and *g_2_* together, i.e., 1 df tests (*g_1_*=0 or *g_2_*=0), or a 2 df test (*g_1_*=*g_2_*=0). The 1 df tests are the tests of main interest, because they distinguish unidirectional from bidirectional causation. Note that the 2 df test is also of interest as the full background confounding (*ra, rc*, and *re* are all non-zero), implies that Ph1 and Ph2 may correlate, regardless of causal relations. We investigated the power in a factorial design, in which each parameter featured as a factor, with a given number of levels.

Additionally, we calculated the R^2^ (proportion of explained variance) in the regression of the phenotypes (Ph1 and Ph2 in Figure 2) on their respective PSs, and the correlation between Ph1 and Ph2 (Table 4). Given the NCPs and the parameters, we proceeded by regressing the NCPs on the parameter values to gauge the influence of the parameter values on the NCPs (i.e, the statistical power). In this manner, we can assess which parameters, in addition to the parameters g_1_ and g_2_, are the most relevant in determining the statistical power. Note that the regression analyses estimate the effect of the parameter values on the NCPs. So they do not directly address statistical power, which depends on the sample sizes, effect sizes, and *α* error rate. To provide some insight into the power given N=1000 MZ and N=1000 DZ, we present power and *R*^2^ for a variety of parameter configurations in Tables 2,3, 4 and S1.

**Table 4.** Exemplary *b_1_, b_3_, g_1_* and *g_2_* values and the power for estimating *g_1_*. Based from the same factorial AE design presented in Table 3 and parameters set presented in Table 1. Highest power with extreme positive *b_1_, b_3_*, and *g_1_* values. Lowest power with lowest g_1_ and g_2_ values. Ncpg_1_, non-centrality parameter for rejecting g_1_ = 0;powg_1_, power for rejecting g_1_ = 0; Ph2_Ph1, *R^**2**^* of the regression of Ph2 on Ph1; PS1_Ph1, R^2^ of the regression of Ph1 on PS1; PS2_Ph2, *R^**2**^* of the regression of Ph2 on PS2.

| $b_1$ | $b_3$ | $g_1$ | $g_2$ | $rf$ | Ph2_Ph<br>1 | PS1_Ph1 | PS2_Ph2 | ncp$g_1$ | pow$g_1$ |
| --- | --- | --- | --- | --- | --- | --- | --- | --- | --- |
| 0.07<br>5 | 0.07<br>5 | 0.0<br>6 | 0.06 | 0.00 | 0.214 | 0.066 | 0.066 | 15.97 | 0.979 |
| 0.07<br>5 | 0.02<br>5 | 0.0<br>6 | 0.02 | 0.00 | 0.140 | 0.068 | 0.023 | 15.97 | 0.979 |
| 0.07<br>5 | 0.07<br>5 | 0.0<br>6 | 0.02 | 0.00 | 0.138 | 0.068 | 0.066 | 15.97 | 0.979 |
| 0.07<br>5 | 0.05<br>0 | 0.0<br>6 | 0.04 | 0.00 | 0.180 | 0.067 | 0.045 | 15.97 | 0.979 |
| 0.07<br>5 | 0.07<br>5 | 0.0<br>6 | 0.04 | 0.00 | 0.180 | 0.067 | 0.066 | 15.97 | 0.979 |
| 0.02<br>5 | 0.05<br>0 | 0.0<br>6 | 0.02 | 0.00 | 0.334 | 0.022 | 0.040 | 4.71 | 0.583 |
| 0.02<br>5 | 0.07<br>5 | 0.0<br>6 | 0.06 | 0.00 | 0.417 | 0.020 | 0.059 | 4.71 | 0.583 |
| 0.07<br>5 | 0.02<br>5 | 0.0<br>2 | 0.02 | 0.25 | 0.126 | 0.070 | 0.026 | 4.70 | 0.582 |
| 0.02<br>5 | 0.05<br>0 | 0.0<br>2 | 0.02 | 0.50 | 0.465 | 0.025 | 0.045 | 1.13 | 0.186 |
| 0.02<br>5 | 0.05<br>0 | 0.0<br>2 | 0.04 | 0.50 | 0.508 | 0.026 | 0.045 | 1.13 | 0.186 |
| 0.02<br>5 | 0.07<br>5 | 0.0<br>2 | 0.02 | 0.50 | 0.463 | 0.026 | 0.065 | 1.13 | 0.186 |
| 0.02<br>5 | 0.02<br>5 | 0.0<br>2 | 0.06 | 0.50 | 0.539 | 0.024 | 0.024 | 1.13 | 0.186 |
| 0.02<br>5 | 0.07<br>5 | 0.0<br>2 | 0.04 | 0.50 | 0.506 | 0.027 | 0.065 | 1.13 | 0.186 |

We executed three factorial designs, i.e., combining all chosen parameters’ values. The first factorial design included full ACE confounding. Each parameter in the parameter vector θ (except *x, y, e_1_*, and *e_2_*) was a factor (a total of 13 parameters), each factor had two levels (Table 1), *e_1_* was specified as 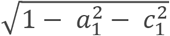 and *e_2_* as 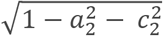. Therefore, the final number of cells in the first design was 2^12^=4096 (Table 1 and 2).

The second design was based on an AE model, i.e., we set *rc=c_1_*=*c_2_*=0. The model without C is of interest, because in adults C influences are often negligible or absent (Polderman et al., 2015). The parameters (i.e., factors) had two or three levels, again *e_1_* was specified as 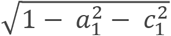 and *e2* as 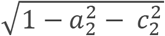. (Table 1 and 3). The number of cells in this design was 3^7^*2^2^= 8748. Table 4 contains example cells of this design, where the power to reject *g_1_* = 0 (df = 1) was moderate to high.

We executed a third design, again based on an AE model. In this design, we included negative and positive *g_1_* and *g_2_* values, which may be of interest given hypothetical negative causal effects (Table 5). In this design, the factors had one or three levels, *e_1_* was specified as 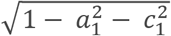 and *e2* as 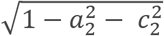, and the number of cells in the first design was 4^4^=256 (Table 1 and 4). Although we focused on the df = 1 tests throughout, we also report the df = 2 tests in Table S1 for this design.

**Table 5.**
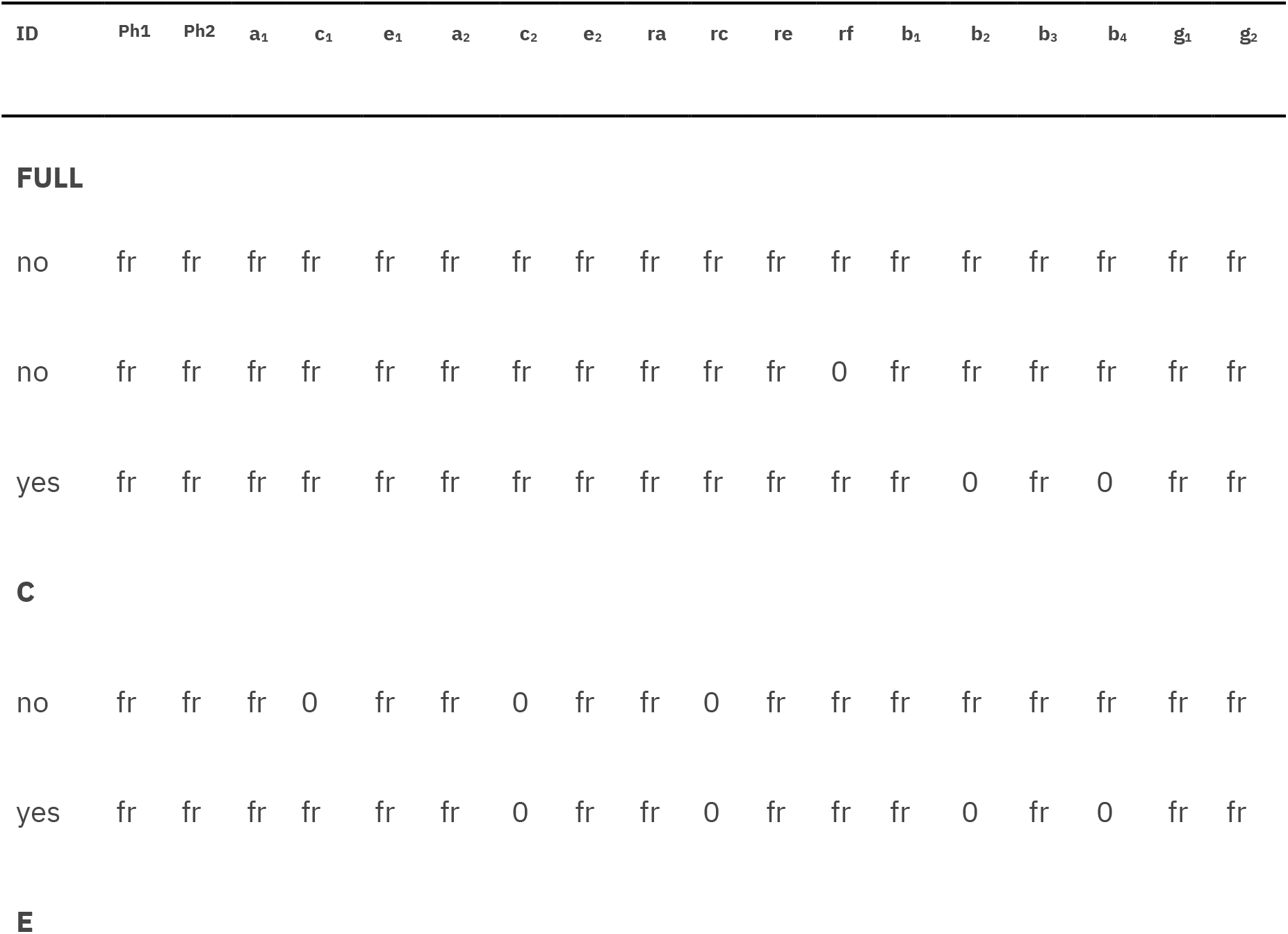
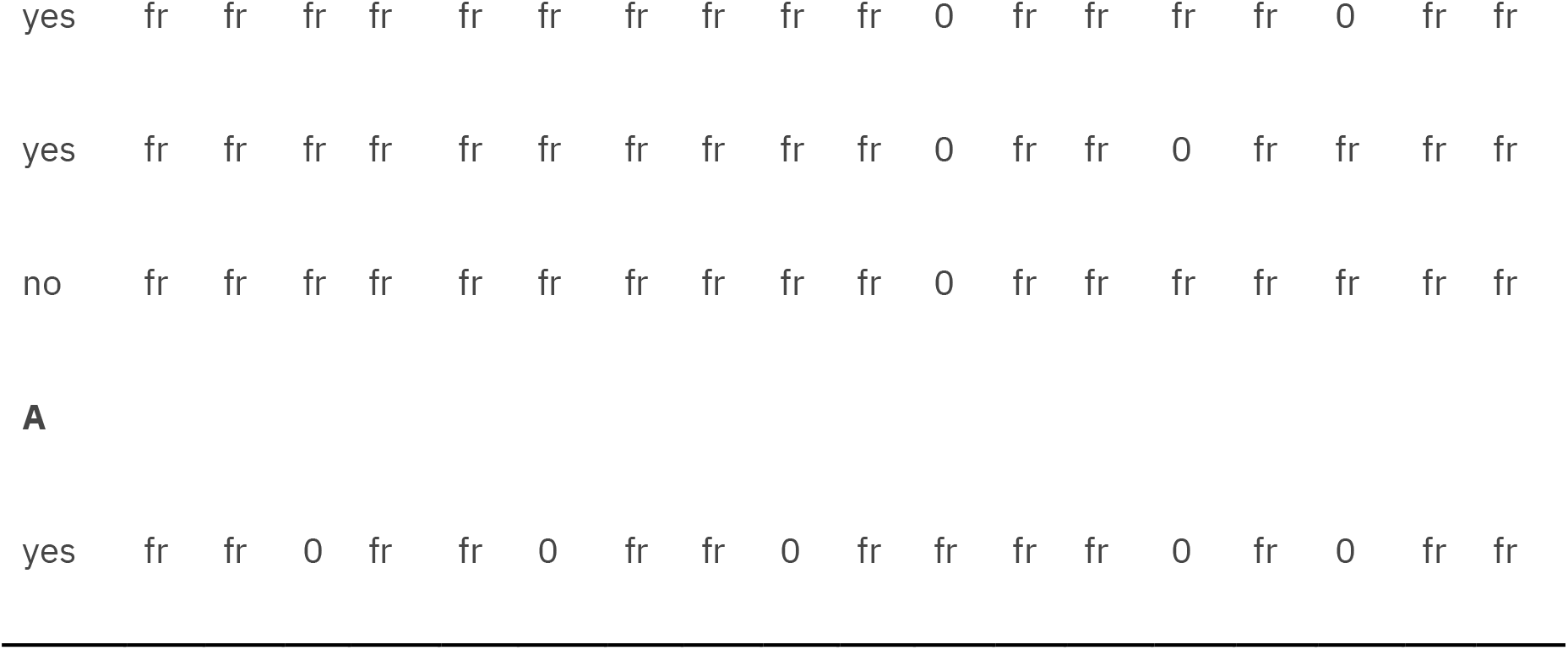
Model Identification. Configurations of fixed and free parameters that may identify the model in Figure 2. Each row shows a combination of fixed or free status for the parameters; the ID column indicates whether the model is locally identified. The text ‘fr’ indicates that the parameter is freely estimated, and ‘0’ denotes that the parameter is fixed to zero.

These factorial designs served to determine the contributions of the various parameters to the statistical power to reject *g_1_*=0, *g_2_*=0, and *g_1_*=*g_2_*=0. We selected parameter values such that the phenotypic twin correlations were reasonable (see Table 5) and the predictive strength of the instrument was plausible (Tables 4 and S1). Specifically, we found the highest power with positive *b_1_, b_3_, g_1_* and *g_2_* values and the lowest with lowest *g_1_* and *g_2_* values. Tables 2 and 3 show results of linear regression analyses in which we regressed the NCP on the parameter values. The *R*^2^ of each parameter reveals much of the variance in the statistical power is explained by that parameter. Thus, the contribution of each parameter to the statistical power changes can be quantified and compared.

## Results

### Power

Based on the simulation results, we identified the following trends concerning the power to reject *g_1_*=0, *g_2_*=0 and *g_1_*=*g_2_*=0. First, the magnitude of the parameters *b_1_* and *b_3_*, i.e., the predictive strength of the genetic instruments, are important to the power. This result can be seen in Tables 2 and 3 by considering the R^2^ values of *b_1_* and *b_3_*. Specifically, the PS regression coefficients, *b_1_* and *b_3_*, explain around 46% of the variance in the NCP. The causal parameters *g_1_* and *g_2_* explain around 40% of the variance in the NCP in the AE models, Table 3. In practice, we found high power (> 0.8, alpha = 0.05, Nmz=Ndz=1000) in a variety of situations with a range of positive and negative values for *b_1_, b_3_, g_1_*, and *g_2_* (Tables 4 and S1).

Based on design 2, we found that the correlation between the instruments (*rf*, Figure 2) made little difference to the power, as is clear from the *R^2^* in Tables 2 and 3. Also, we found that the background correlations, i.e., the *ra, rc* and *re* parameters, are relatively unimportant to the power in the present designs (Table 2 and 3).

We performed two factorial design analyses (with and without C variance), because we wanted to assess how the presence of C variance would affect the power. In Table 2, which includes C variance, *b_1_* and *b_3_* explain 36% of the variance of the NCP, which is less than the variance explained in the absence of C variance in the model (namely, 46% R^2^ in the AE model, Table 2). So, while we see that the parameters related to C have little influence on the power, the absence of C is beneficial. Needless to say, whether C can be omitted in practice will be dictated by the data.

We found that differences in the variance components of *Ph1* and *Ph2* (i.e., differences in the parameters *a_1_* vs. *a_2_* and *c_1_* vs. *c_2_*) had little effect on the power. This result is at odds with traditional DoC models, in which the statistical power depends heavily on these differences (Gillespie et al., 2003). Strikingly, in the present model, differences between the phenotypes in variance decomposition appear unimportant to the statistical power.

The present results were based on a linear regression of the NCPs on the parameters. We investigated possible non-linearity by including second order polynomials of the parameters. However, we found that the second order terms explained 7.1-8.9% of the variance in the NCPs (g1=0, Tables 2 and 3). From tables 2 and 3 the R^2^ ranged from 90.2% (NCP regressed on all other parameters) in the AE model to 94.5% in the ACE model.

In an attempt to put the power analyses in perspective to published results, we present the power curves for four realistic scenarios (Figure 3). The factorial design was performed with values for additive and shared genetic variances based on previous work (see below). The number of twin pairs required to achieve an 80% power of rejecting g1 = 0 is plotted against an increasing R^2^ for the instrument PS1 (path b_1_). Environmental variances were dependent on shared and additive genetic variances so that *e_1_* was specified as 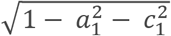 and *e2* as. 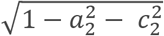 Additive and shared variances were considered as follows: (A) alcohol use (*a^2^* 49%, *c^2^* 10%) (Verhulst et al., 2015) and heart disease (*a^2^* 22%, *c^2^* 0%) (Wu et al., 2014); (B) BMI (*a^2^* 72%, *c^2^* 3%) (Rokholm et al., 2011) and major depression (*a^2^* 37%, *c^2^* 1%) (Scherrer et al., 2003); (C) cannabis use (*a^2^* 51%, *c^2^* 20%) (Verweij et al., 2010) and schizophrenia (*a^2^* 81%, *c^2^* 11%) (Sullivan et al., 2003); (D) dyslipidemia (LDL) (*a^2^* 60%, *c^2^* 28%) (Zhang et al., 2010) and again heart disease (*a^2^* 22%, *c^2^* 0%) (Wu et al., 2014) (Figure 3). Figure 3 also includes vertical lines with previously measured estimations of four PSs: Smoking **(Pasman et al., 2022)**, BMI (Furlong and **Klimentidis, 2020)**, LDL (Kuchenbaecker et al., 2019), and attention deficit hyperactivity disorder (ADHD) (Demontis et al., 2019). Figure 3 shows that the power to reject *g1*=0 is quite reasonable and well within values that appeared in recent publications.

**Figure 3.**
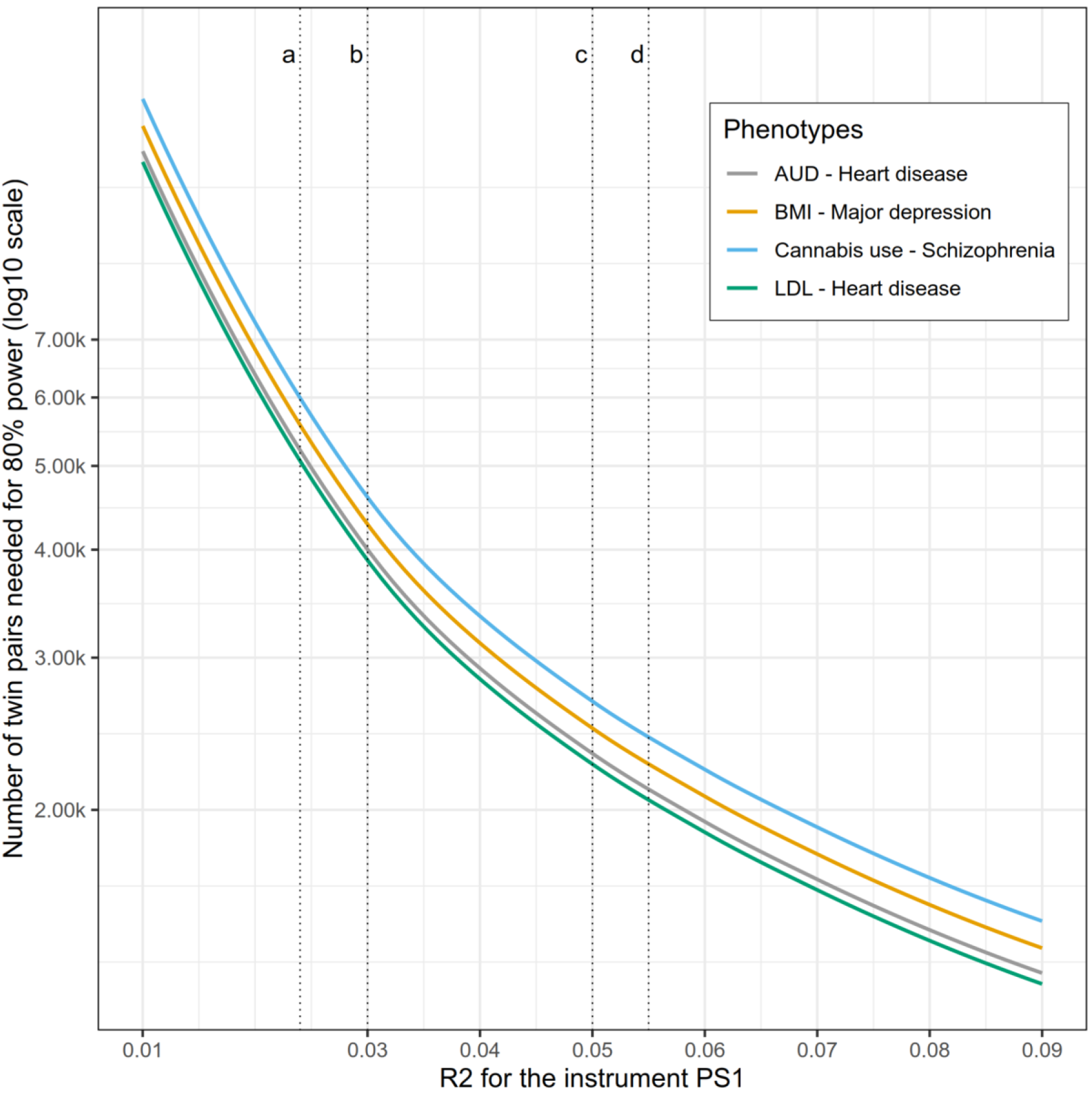
Power curve across R^2^ values for PS1. Notes: Parameters set for all groups 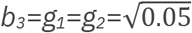; *ra*=.3; *rc*=.25; *re*=.3; *rf*=.25; *re* =.3; *rf* =.25; Environmental variances were dependent on shared and additive genetic variances: *e_1_* was specified as 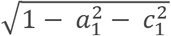 and *e_2_* as 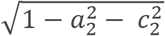. Additive and shared variances for the groups: (A) cannabis use (a^2^ 51%, c^2^ 20%) (Verweij et al., 2010) and schizophrenia (*a^2^* 81%, *c^2^* 11%) (Sullivan et al., 2003); (B) BMI (*a^2^* 72%, *c^2^* 3%) (Rokholm et al., 2011) and major depression (*a^2^* 37%, *c^2^* 1%) (Scherrer et al., 2003); (C) alcohol use (*a^2^* 49%, *c^2^* 10%) (Verhulst et al., 2015) and heart disease (*α^2^* 22%, *c^2^* 0%) (Wu et al., 2014); (D) dyslipidemia (LDL) (*a^2^* 60%, *c^2^* 28%) (Zhang et al., 2010) and heart disease (*a^2^* 22%, *c^2^* 0%) (Wu et al., 2014). Vertical lines were added to represent R^2^ for four PSs reported in recent papers: a, smoking (**Pasman et al., 2022**); b, BMI (**Furlong and Klimentidis, 2020**); c, LDL (Kuchenbaecker et al., 2019); d, attention deficit hyperactivity disorder (ADHD) (Demontis et al., 2019).

### Discussion

We presented a new twin model based on the MR-DoC twin model (Minica, et al., 2018), which we refer to as the MR-DoC2 model. Compared to the MR-DoC model, MR-DoC2 model accommodates full confounding (originating in background A, C, and E effects) and bidirectional causality. The model can be used to investigate the bidirectional causal interrelationship. As usual in twin studies, the 2×2 phenotypic covariance matrix was decomposed into A, C, and E covariance matrices. However, in addition, phenotypic reciprocal causal paths were specified, as well as paths from the PSs to phenotypes Ph1 and Ph2.

The MR-DoC2 is suitable for the study of bidirectional causality given A, C, and E confounding, whilst retaining reasonable statistical power. While we have assumed no direct effects of PS1 on Ph2 and PS2 on Ph1, PS1 is correlated with Ph2 and PS2 is correlated with Ph1. Given non-zero *rf*, the correlation between PS1 and PS2, PS1 is correlated with Ph2 via the *rf* path (see Figure 2). The model is based on the assumption that there are no direct effects of the PS1 on the outcome Ph2 (and PS2 on Ph1), but there are indirect associations with Ph2 from PS1 via *rf* when *rf* does not equal zero. Pleiotropy also occurs though the causal feedback loop if both *g_1_* and *g_2_ are non-zero*. As shown in Table 5, there are situations in which the paths b_2_ and b_4_ can be estimated, but these involve other constraints. For instance, either b_2_ or b_4_ (but not both) can be estimated if the unshared environmental correlation, *re*, is fixed to zero.

As expected, statistical power is higher when a larger proportion of phenotypic variance is explained by the genetic instruments, PS1 and PS2. This is encouraging, because the proportion of trait variance accounted for by SNPs is likely to increase with greater sample sizes and with improvements in the ability to test rare variants’ associations with outcomes and their risk factors. We found the sign of the causal parameters (*g_1_, g_2_*) to be relevant to the power to reject *g_1_*=0 or *g_2_*=0. Negative values (*g_1_*<0, *g_2_*<0) or a combination of negative and positive values were associated with high power (Table S1). This result is consistent with one previously reported in multivariate twin studies (Evans & Duffy, 2004), where power increases with negative genetic correlation.

In cross-sectional DoC modeling, the statistical power to discern direction of causation is greater when the phenotypes have different variance component proportions. That is to say, if the MZ and DZ correlations for exposure equal those for the outcome variable, the model will not be able to detect causal direction (Heath et al., 1993). However, in the present article we have shown that the addition of the PSs permit a larger range of genetic decomposition scenarios and these components (a_1_, c_1_, e_1_, a_2_, c_2_, e_2_) have little effect on the ability to detect causal directionality, which is a remarkable advantage over the original DoC model. More importantly, this model accommodates A, C, and E confounding, improving on previously noted limitations of DoC models (Rasmussen et al., 2019).

We presented a model in which the background covariance structure involved the same sources of variances, i.e., ACE or AE. It may happen that the phenotypes differ in this respect (e.g., ACE and ADE). This poses no problem with respect to model specification. This situation has the advantage of allowing b2 or b4 to be estimated, because the background confounding is necessarily limited to A and E (i.e, ra and re). We did not pursue this here as the combination of ACE and ADE phenotypes seems somewhat uncommon, in part because of the sample size.

Another scenario that is worth noting is when one considers a PS as an imperfect measure of the additive genetic liability, where *rf* asymptotically tends to *ra*. We did establish that a constraint *ra=m*rf*, with fixed m (e.g, m=1), identified b2 or b4 (direct pleiotropic paths), but not both. If the relationship of *ra* and *rf* is of interest, a sensitivity analysis with varied values of the constant *m* can be set up to evaluate the effect on the causal estimates. In the case *m*=1 it is implied that *ra=rf*.

We note that in Tables 2 and 3, the R^2^ values did not sum to unity. However, the variances for *b_1_, b_3_, g_1_*, and *g_2_* account for over 88% of the NCP variance, while the remainder are due to non-linear effects and interactions. We consider these to be too situation-specific and, therefore, of little interest in conducting a power analysis. All results are based on simulations of 1000 MZs and 1000 DZs, however, the code is available in a repository (https://github.com/lf-araujo/mr-doc2), so the reader can perform their own power calculations and include the number of observations that best suits their study.

In the standard DoC model, parameter estimates can be biased if the reliabilities of the two variables differ (Heath et al., 1993). Specifically, the bias in the causal parameters is towards the more reliable variable being the cause of the less reliable one. In additional simulations, we established that unmodeled measurement error (phenotypic reliabilities < 1) resulted in bias in the estimates of the parameters e_1_, e_2_, and re. While the power to reject g_1_=0 and/or g_2_=0 was lower in the presence of measurement error, the actual estimates of the causal parameters *g_1_* and *g_2_*, and the estimates of the parameters *b_1_* and *b_3_* were unaffected by unmodeled measurement error.

As seems inevitable when trying to establish causation in non-experimental settings, some assumptions are necessary. It is important to emphasize that although the model incorporates (indirect) horizontal pleiotropy through *rf*b3* or through *rf*b1*, the absence of the direct horizontal pleiotropy is a required assumption (i.e., the parameters b2 and b4 are zero). It represents a portion of the total horizontal pleiotropy (we assume that the total horizontal pleiotropy consists of *rf*b3* + *rf*b1* + *b2* + *b4*). Violation of this assumption (*b2* and *b4* not equal 0) results in bias in the causal parameters g1 and g2 and in the parameters ra, rc, and re. Specifically, given b_2_ > 0 and b_4_ > 0, the parameters g_1_ and g_2_ are overestimated as a consequence of fixing b_2_ = b_4_ = 0. This overestimation increases the false positive rate, i.e., the rate with which we incorrectly reject g_1_=0 and/or g_2_=0. Given b_2_ > 0 and b_4_ > 0, the parameters ra, rc, and re are underestimated, as a consequence of the overestimation of g_1_ and g_2_. Furthermore, a mismatch between the phenotype in the GWAS that generated the PS and the phenotype used in MR-DOC2 could alter estimates. As the mismatch between the phenotypes in the discovery GWAS used to create the PRS and the MR-DoC2 phenotype increases, the b1 and b3 paths will become smaller, become weaker instruments, and potentially reduce power. The size of this effect was not pursued in this paper, and will depend on the nature of the mismatch.

The use of instrumental PS is common in Mendelian randomization studies (Burgess et al., 2020; Dudbridge, 2021). It has been shown that the use of a PS as an instrument is mathematically equivalent to a weighted mean of the results from individual SNPs (Dudbridge, 2021). However, its use also comes with challenges. It plays the role of a stronger instrument, but it is also a composite of variants that may themselves have indirect or direct effects on the outcome. Therefore, using a PS increases the risk of (horizontal) pleiotropy when compared to the use of a single variant. Note that the present model does account for several types of pleiotropy already in the model. First, in Figure 2, the parameter *rf* represents the correlation between the two PSs, which may partially account for a correlation between PS1 (the instrument of *Ph1*) and *Ph2*. Second, the bidirectional causal paths (parameters between *Ph1* and *Ph2* also connect PS1 and Ph2 and PS2 and Ph1. A useful feature of the MR-DoC2 model is the inclusion of non-shared (E) confounding (parameter re). The constraint re=0 is considered a drawback of standard DoC models (Rasmussen et al., 2019), which also applied to the MRDoc model, as presented by Minica et al. (2018).

Multivariate, GxE and DoC approaches have been considered less practical because they require larger data sets to obtain accurate estimates of parameters relating two traits (Gillespie et al., 2003). More recently, however, the availability of studies with very many participants have made these models practical to apply. We have shown that the MR- DoC2 model has moderate power to test bidirectional causation, and therefore is suitable for a range of clinical applications. The next steps in model investigation will include extensions for longitudinal and multivariate data, to provide corroboration of potential causal pathways identified by this modeling.

## Declarations

## Ethics approval

Not applicable

## Consent for publication

Not applicable

## Availability of data and material

Data sharing not applicable to this article as no datasets were generated or analyzed during the current study.

## Code availability

Code is available in a repository for replication.

---

**Table S1.**
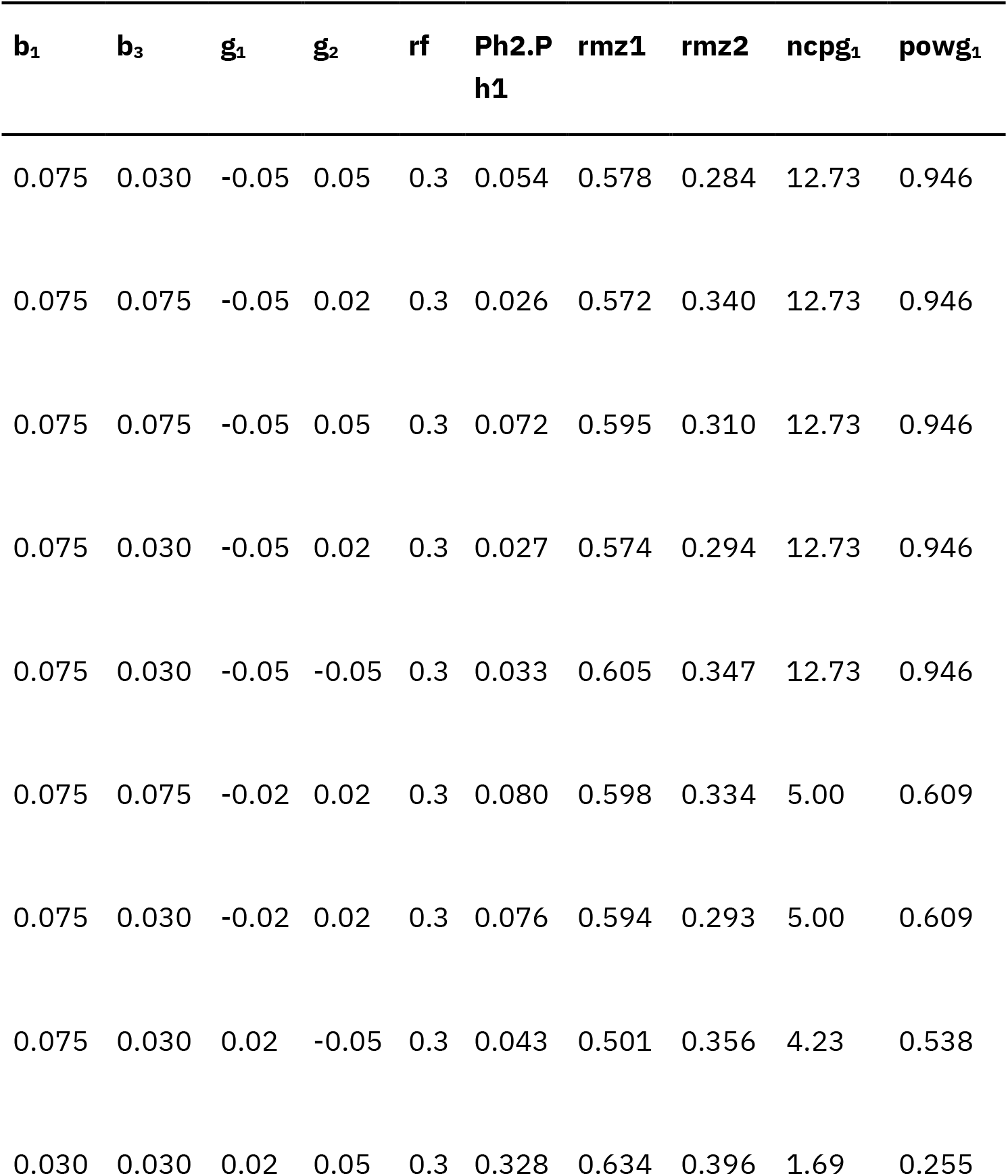
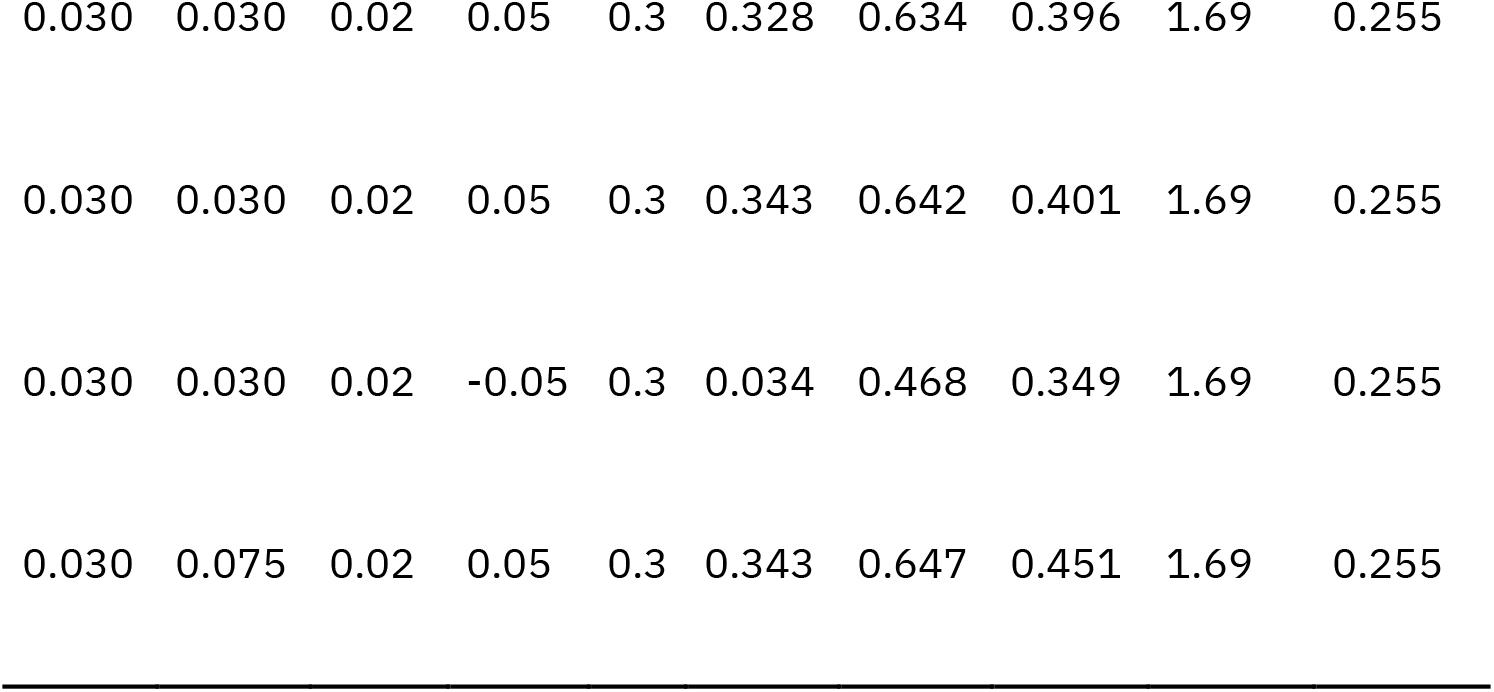
Exemplary *b_1_, b_3_, g_2_* and *g_2_* values and the power for estimating g1. High confounding (*ra*=*re*=.3), asymmetric AE (*a_1_*± =.5, *e_1_* =.5, *a_2_* =.3, *e_2_*=.7). Highest power with combinations of positive b_1_, b_3_, with negative g_1_ or g_2_ values. Ncpg1, noncentrality parameter for rejecting g1 = 0;powg_1_, power for rejecting g_1_ = 0; Ph2.Ph1, *ß^**2**^* of the regression of Ph2 on Ph1. Rmz1 and 2, phenotypic twin correlations.

